# Label-free identification of non-activated lymphocytes using three-dimensional refractive index tomography and machine learning

**DOI:** 10.1101/107805

**Authors:** Jonghee Yoon, YoungJu Jo, Min-hyeok Kim, Kyoohyun Kim, SangYun Lee, Suk-Jo Kang, YongKeun Park

**Affiliations:** Department of Physics, Korea Advanced Institute of Science and Technology (KAIST), Daejeon 34141, South Korea; KAIST Institute Health Science and Technology, Daejeon 34141, South Korea; Department of Biological Sciences, KAIST, Daejeon 34141, South Korea; Tomocube, Inc., Daejeon 34051, South Korea

**Author notes:** These authors contributed equally to this work (Jonghee Yoon, YoungJu Jo). Correspondence: YongKeun Park, PhD, KAIST, 291 Daehak-ro, Yuseong-gu, Daejeon 305-701, Korea.

## Abstract

Identification of lymphocyte cell types is crucial for understanding their pathophysiologic roles in human diseases. Current methods for discriminating lymphocyte cell types primarily relies on labelling techniques with magnetic beads or fluorescence agents, which take time and have costs for sample preparation and may also have a potential risk of altering cellular functions. Here, we present label-free identification of non-activated lymphocyte subtypes using refractive index tomography. From the measurements of three-dimensional refractive index maps of individual lymphocytes, the morphological and biochemical properties of the lymphocytes are quantitatively retrieved. Machine learning methods establish an optimized classification model using the retrieved quantitative characteristics of the lymphocytes to identify lymphocyte subtypes at the individual cell level. We show that our approach enables label-free identification of three lymphocyte cell types (B, CD4+ T, and CD8+ T lymphocytes) with high specificity and sensitivity. The present method will be a versatile tool for investigating the pathophysiological roles of lymphocytes in various diseases including cancers, autoimmune diseases, and virus infections.

---

Lymphocytes consisting of various cell types including B, helper (CD4+) T, cytotoxic (CD8+) T, and regulatory T lymphocytes, and play crucial roles in the adaptive immune system^1^. Each lymphocyte cell type has different functions: B lymphocytes produce antibodies, and T lymphocytes recognize a specific antigen and execute effector functions. The lymphocyte population and function are tightly regulated to defend the host against harmful invaders or abnormal conditions^1,2^. Disturbances in lymphocyte function and regulation are related to various diseases including cancers^3-5^, autoimmune diseases^6,7^, and virus infections^8,9^.

To understand the roles of different types of lymphocytes, several methods based on labelling techniques have been developed to identify and discriminate lymphocyte cell types. Because different kinds of lymphocytes have very similar cellular morphology such as a large nucleus with small cytosolic regions and round shapes, conventional optical methods such as bright-field microscopy or phase contrast microscopy have limited ability in classifying lymphocyte cell types^10^. To overcome this, a specific surface membrane proteins, known as surface markers, are recognized and tagged with magnetic beads or fluorescence molecules via antigen-antibody binding. And then, specific types of lymphocytes are identified and separated by magnetic forces or fluorescence signals^11^. Targeting surface markers is a precise and efficient manner to determine the cell types; however, labelling methods have potential risks of altering cellular functions by modifying membrane protein structures. Moreover, labelling methods have limitations such as they cannot simultaneously identify multiple cell types due to the limited numbers of distinguishable labelling agents^12^.

Label-free approaches such as mass spectroscopy^12^ and Raman spectroscopy^10^ have also been introduced to overcome the limitations of labelling methods because these spectroscopic methods exploit intrinsic biochemical properties of lymphocytes. Mass spectroscopy measures cellular biochemical properties which enable the profiling of lymphocyte proteins as well as the identification of lymphocyte subtypes. However, it has a limitation in live-cell analysis due to the homogenization process of the cells. Raman spectroscopy measures molecular vibrations and characterizes biochemical properties of a sample. Raman spectroscopy permits label-free live-cell analysis of lymphocytes with high accuracy; however, it requires a bulky optical system and long acquisition time (several seconds) to measure 2-D Raman signals, which limits its broad use in clinics.

Here, we present a label-free method to identify lymphocyte cell types by exploiting optical diffraction tomography (ODT) and machine learning. ODT is a label-free imaging technique that measures a three-dimension (3-D) refractive index (RI) tomogram of a sample which quantitatively provides morphological and biochemical information^13,14^. Previously, ODT has been widely used to study various biological samples including red blood cells^15-22^, white blood cells (WBC)^23,24^, hepatocytes^25^, cancer cells^16,26-32^, neurons^32,33^, bacteria^34,35^, phytoplankton^36^, and hair^37^. In our previous study, we reported that ODT enables the quantitative analysis of WBCs including lymphocytes and macrophages^23^. We demonstrated that lymphocytes and macrophages could be discriminated based on quantitative morphological and biochemical properties; however, we were unable to identify lymphocyte cell types due to their indistinguishable cellular morphology and biochemical characteristics.

In the present study, to find morphological and biochemical differences in indistinguishable lymphocyte cell types, we used machine learning techniques. Machine learning methods construct classification models by combining multiple features in a data-driven manner, not a hypothesis-driven manner. This approach is especially powerful for high-dimensional data that are extremely difficult to manually process by a human due to the complicity and large size^38^. In biomedical research fields, machine learning methods have been widely used to solve complex biological problems: identification of bacterial species^39^, discrimination of WBC types^40,41^, and investigation of pathophysiologic conditions^42-44^. However, there has been no applications that use both 3-D RI tomography and machine learning for the purpose of biomedical studies. Here, we exploit machine learning techniques to establish classification models using the quantitative morphological and biochemical information of lymphocytes which is retrieved from the 3-D RI tomograms of individual cells and show that established classification models enable the identification of three lymphocyte subtypes (B, CD4+ T, and CD8+ T lymphocytes). The present approach is a versatile tool for determining lymphocyte cell types in a label-free manner.

## Results

The overall procedure for label-free identification of lymphocytes is summarised in Fig. 1. The present method involves three steps: (i) measurement of the 3-D RI tomograms of individual lymphocytes (Fig. 1a), (ii) establishment of a statistical classification model using the quantitative biochemical and morphological features of the lymphocytes obtained from the 3-D RI tomograms (Fig. 1b), and (iii) identification of the lymphocyte types using the established classifier at a single cell level (Fig. 1c).

**Figure 1.**
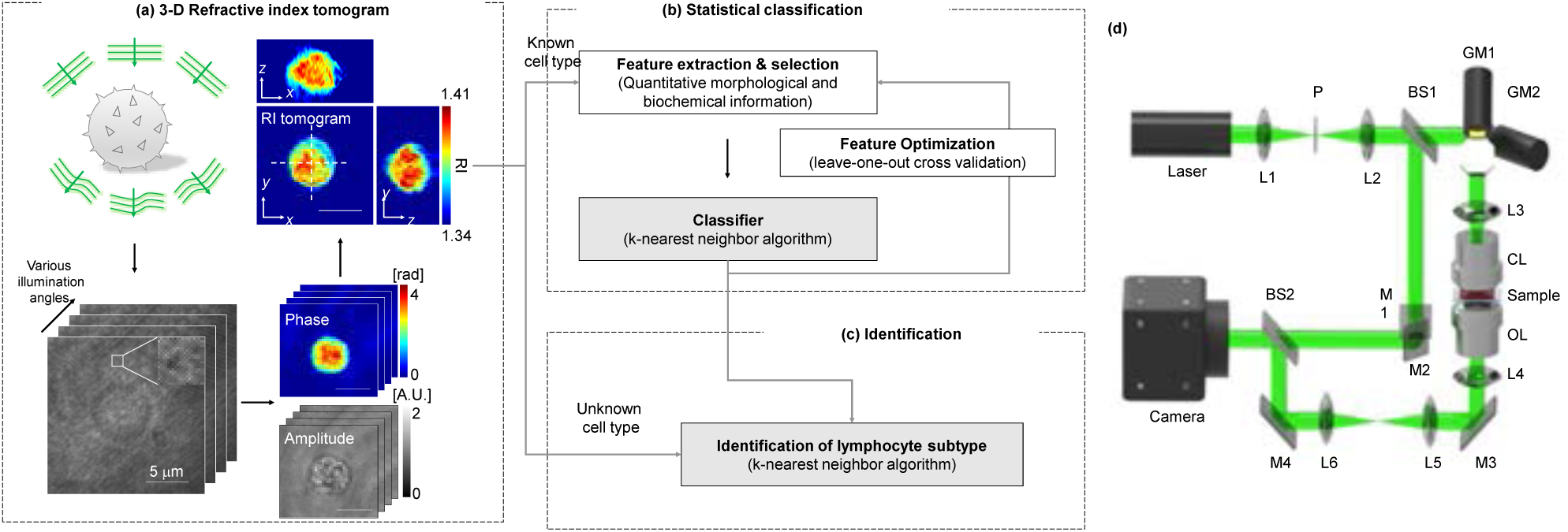
Schematic diagrams of the label-free identification of lymphocyte cell types using optical diffraction tomography and machine learning algorithm. (a) Procedures for measuring label-free 3-D tomograms of lymphocytes. Multiple holograms of lymphocytes are measured by changing the angle of illumination. Optical field information of the lymphocytes was retrieved from the measured holograms, and then, 3-D refractive index tomograms were reconstructed using the retrieved optical field information. Scale bar is 5 μm. (b) Establishing a statistical classifier for identifying lymphocyte cell types using *k*-nearest neighbour algorithm. Quantitative morphological and biochemical features of lymphocytes known their cell types (supervise learning) were used to generate the classifier. (c) Determining the types of unidentified cells with the established classifier. (d) Schematic of the experimental setup. A Mach-Zehnder interferometric microscope equipped with a 2-D scanning galvanomirror (GM) was used for measuring the holograms of lymphocytes. BS1–2, beam splitters; L1–6, lens; SF, spatial filtering; CL, condenser lens; OL, objective lens; M1–4, mirrors.

Figure 1a shows the procedures for reconstructing the 3-D RI tomograms of the lymphocytes. To reconstruct the 3-D RI tomograms, multiple 2-D holograms of a cell are measured at various angles of illuminations using an interferometric microscope (Fig. 1d). A coherent laser beam is split into two arms by a beam splitter. One arm passes through a sample, and then, the diffracted light from the sample is projected onto a camera plane with a microscope. At the camera plane, the sample beam interferes with the other arm and generates a spatially modulated hologram. The angle of the beam impinging onto the sample is controlled by a dual-axis galvanomirror. From the measured holograms, complex optical fields consisting of both the amplitude and quantitative phase images are retrieved using a field retrieval algorithm (See Methods). Then, a 3-D RI tomogram of a lymphocyte is reconstructed using the retrieved multiple optical amplitude and phase information via an optical diffraction tomography algorithm^14,45^.

To test whether 3-D RI tomograms of lymphocytes enable the identification of their cell types, three types of lymphocytes (B cell, CD4+ T, and CD8+ T lymphocytes) were obtained from mice peripheral blood through specific surface marker staining and flow cytometry (See Methods) prior to measuring on the 3-D RI tomograms. Figure 2 shows representative 3-D RI tomograms of each lymphocyte type. Cross-sectional slices of the representative 3-D RI tomograms of B cell, CD4+ T cell, and CD8+ T cell in the *x*-*y*, *y*-*z*, and *x*-*z* planes are also shown in Figs. 2a-c, respectively. The measured RI distribution clearly represents the cellular boundaries and internal organelles such as the nuclear membrane and nucleoli. The B cell shows a well-defined nucleus and nucleoli with RI values ranging from 1.34 to 1.41, while the RI values of the cytosolic areas of CD4+ and CD8+ T cells are higher than that of the B cell. 3-D RI tomograms of each lymphocyte type in Figs. 2a-c are rendered using a customized transfer function in a commercialized software (Tomostudio^TM^, Tomocube Inc., Republic of Korea) to resemble haematoxylin and eosin staining (Figs. 2d-f and Supplementary Videos 1-3). These 3-D rendered images show their round cellular boundaries and intracellular components. Even though each lymphocyte type looks like it has a slightly different RI distribution, they are basically indistinguishable due to their identical cellular size and morphology.

**Figure 2.**
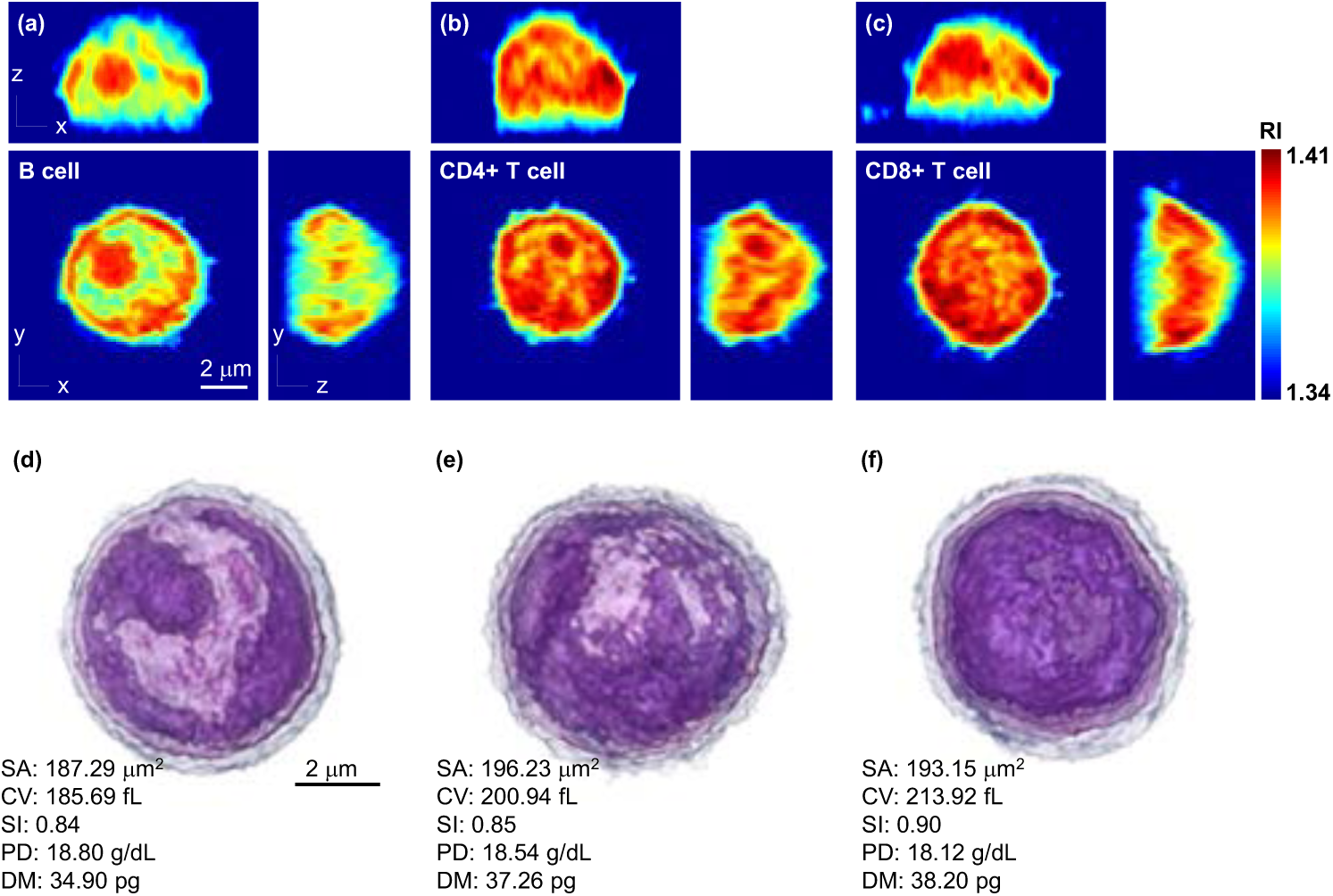
Representative 3-D RI tomograms of each lymphocyte type and the 3-D rendered images with quantitative characterization. Cross-sectional slices of a RI tomogram of (a) B cell, (b) CD4+ T cell, and (c) CD8+ T cell. Scale bar is 2 μm. (d-f) 3-D rendered RI tomograms and quantitative characterization of the morphological and biochemical features of (a-c), respectively. Scale bar is 2 μm. SA, surface area; CV, cellular volume; SI, sphericity; PD, protein density; DM, dry mass.

To investigate the differences among the lymphocyte cell types, a statistical analysis was conducted of the quantitative characteristics of the three lymphocyte subtypes. Quantitative morphological (cellular surface area, cellular volume, and sphericity) and biochemical (cellular dry density and dry mass) parameters were calculated from the 3-D RI distribution of the lymphocytes (See Methods). Figures 2d-f show the quantitative characteristics of the B, CD4+ T, and CD8+ lymphocytes, respectively. The cellular surface area and volume of lymphocytes were simply calculated from the voxel information of the 3-D RI tomograms. And then, the sphericity, a dimensionless parameter that indicates the roundness of the cellular morphology, was obtained by the ratio of the calculated surface area and volume. The biochemical information was retrieved from the RI distribution of the lymphocytes because the RI values are linearly proportional to the local concentration of non-aqueous molecules (mostly protein).

Figure 3 shows scatter plots of the quantitative morphological and biochemical information of each lymphocyte type and the results of the statistical analyses. The cellular surface areas of the B cell (*n* = 149), CD4+ (*n* = 95), and CD8 T lymphocytes (*n* = 111) were 145.87 ± 20.25, 167.23 ± 32.08, and 160.80 ± 19.12 μm^2^, respectively (Fig. 3a). The B cells had significantly smaller cellular surface areas compared to the T cell subtypes (*p*-value < 0.001), while there was no statistical difference between the CD4+ and CD8+ T cells. In addition, the result of quantifying the cellular volume of the lymphocytes also shows a similar tendency with the result of the cellular surface area analysis. The cellular volume of the B lymphocytes (133.43 ± 26.47 fL) was significantly small (*p*-value < 0.001) compared to that of the CD4+ (155.73 ± 35.14 fL) and CD8+ (152.77 ± 26.52 fL) lymphocytes (Fig. 3b). However, the CD4+ and CD8+ T cells had a similar cellular volume. The sphericities were 0.86 ± 0.06, 0.84 ± 0.06, and 0.86 ± 0.05 for the B cells, CD4+ and CD8+ T cells, respectively (Fig. 3c). Although the sphericity of the CD4+ T cells was statistically smaller than that of the B cells (*p*-value < 0.01) and CD8+ T cells (*p*-value < 0.05), all the lymphocyte subtypes had a high sphericity which means the lymphocytes have round shapes.

**Figure 3.**
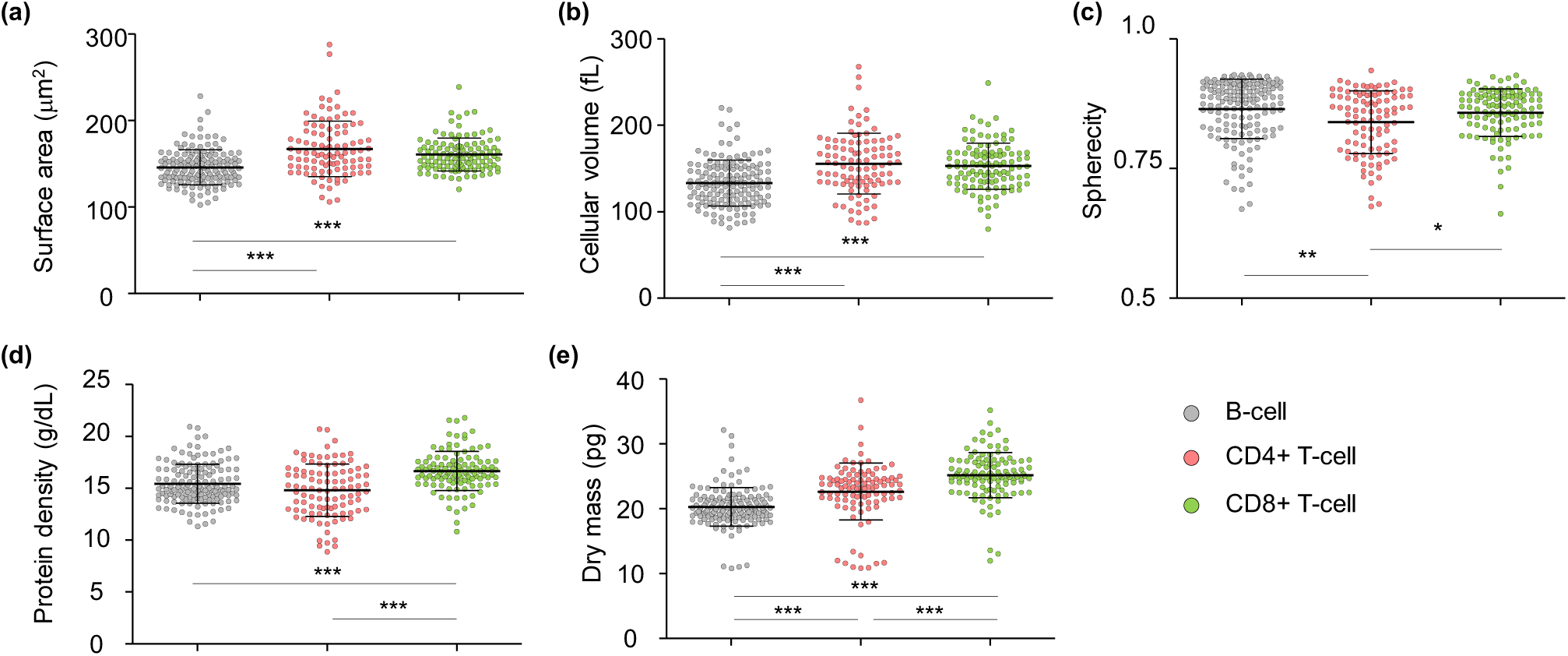
Quantitative analysis of the morphological and biochemical features of the B cell, CD4+ T cell, and CD8+ T cell. (a) Surface area, (b) cell volume, (c) sphericity, (d) protein density, and (e) dry mass. Each symbol indicates an individual cell measurement. Horizontal black lines indicate the mean values; vertical lines, indicating standard deviation. Statistical analysis was done with Student’s *t*–test.

Next, we compared the biochemical properties of the lymphocytes. The cellular dry mass density of the B cells, CD4+ and CD8+ T cells was 15.43 ± 1.88, 14.81 ± 2.54, and 16.66 ± 1.88 g/dL, respectively (Fig. 3d). The CD8+ T cells had a significantly increased cellular dry mass density compared to that of the others (*p*-value < 0.001). The total cellular dry mass, calculated by integrating the dry mass density over the cell volume, was 20.28 ± 2.97, 22.65 ± 4.49, and 25.19 ± 3.51 pg for the B cells, CD4+ and CD8+ T cells, respectively (Fig. 3e). There were significant differences in the total cellular dry mass among the cell types (*p*-values < 0.001). The B cells had a smaller cellular mass compared to the T cell subsets. Moreover, the CD8+ T cells were statistically heavier than that of the CD4+ T cells. These results reveal that the biochemical characteristics of each lymphocyte type have statistical differences. However, even though the quantitative morphological and biochemical analyses show statistical differences among the lymphocyte population, a single lymphocyte cannot be identified by its cell type using a single retrieved quantitative characteristic because of cell-to-cell variations.

To identify the lymphocyte types at an individual cell level, machine learning methods were exploited to construct classification models using multiple quantitative characteristics of the lymphocytes. To demonstrate a proof of concept, a training-and-identification method, called statistical classification or supervised machine learning, was used. Seventy percent of the total lymphocytes were randomly chosen and systematically analysed to extract the unique characteristics for each lymphocyte type (Fig. 1b), and then, the cell types of the remaining lymphocytes were identified with the extracted features (Fig. 1c). For the statistical model, a *k*-NN algorithm was used^46,47^, which has been widely used for classification purposes in various research fields. The *k*-NN algorithm consists of a supervised machine learning which establishes a classifier based on quantitative characteristics of the training lymphocytes from known cell types. In this study, a four nearest neighbour (*k*-NN, *k* = 4) algorithm was used to establish the classification models.

To exploit the structural and biochemical information of the intracellular organelles of the lymphocyte as features of the *k-*NN (*k* = 4) algorithm, the quantitative characteristics of the lymphocytes were calculated at various threshold RI values by increasing the threshold from 1.34 to 1.378 with an increment of 0.002. We found that the 3-D RI distribution, which had higher RI values than the threshold RI, tends to reveal information about the intracellular components as the threshold RI values increase (Supplementary Fig. 1). All combinations of structural and biochemical information obtained at a single RI threshold or two different RI thresholds are tested to establish an optimized classifier with cross-validation, and then, a classifier, which shows the best performance, is selected. Then, the remaining lymphocytes are identified using the established classifier, and the identification accuracy (sensitivity and specificity) is measured.

Figure 4 and Table 1 show the overall cross-validation accuracy, sensitivity (true positive results overall positive inputs), and specificity (true negative results overall negative inputs) of the training and test results. We performed statistical classification on three different combinations of lymphocyte types: (i) B and T lymphocytes, (ii) T lymphocyte subsets (CD4+ and CD8+), and (iii) all three types of lymphocytes. First, the T cell subsets were considered as one T cell type, and the classification model is optimized to have the best overall cross-validation accuracy. Features used for establishing the classifier were the surface area, sphericity, and dry mass (RI threshold: 1.342) and all the quantitative information (RI threshold: 1.368). The overall accuracy of the optimized classifier for identifying the B and T cells was 93.15%, and the test accuracy was 89.81% (Fig. 4a). Second, the CD4+ and CD8+ T cells were analysed. To optimize the statistical classification model, the surface area and sphericity (RI threshold: 1.342 and 1.362) of the CD4+ and CD8+ T cells, respectively, were used as features. The classification accuracy was 87.41% and 84.38% for the training and test, respectively (Fig. 4b). Lastly, the statistical classification was performed to identify the three types of lymphocytes. The classification model was optimized with the surface area, dry mass density (RI threshold: 1.340), and sphericity, dry mass density, dry mass (RI threshold: 1.370) as features. The accuracy of the training and test were 80.65% and 75.93%, respectively. These results indicate that machine learning enables the identification of lymphocyte cell types with an accuracy of over 75%.

**Figure 4.**
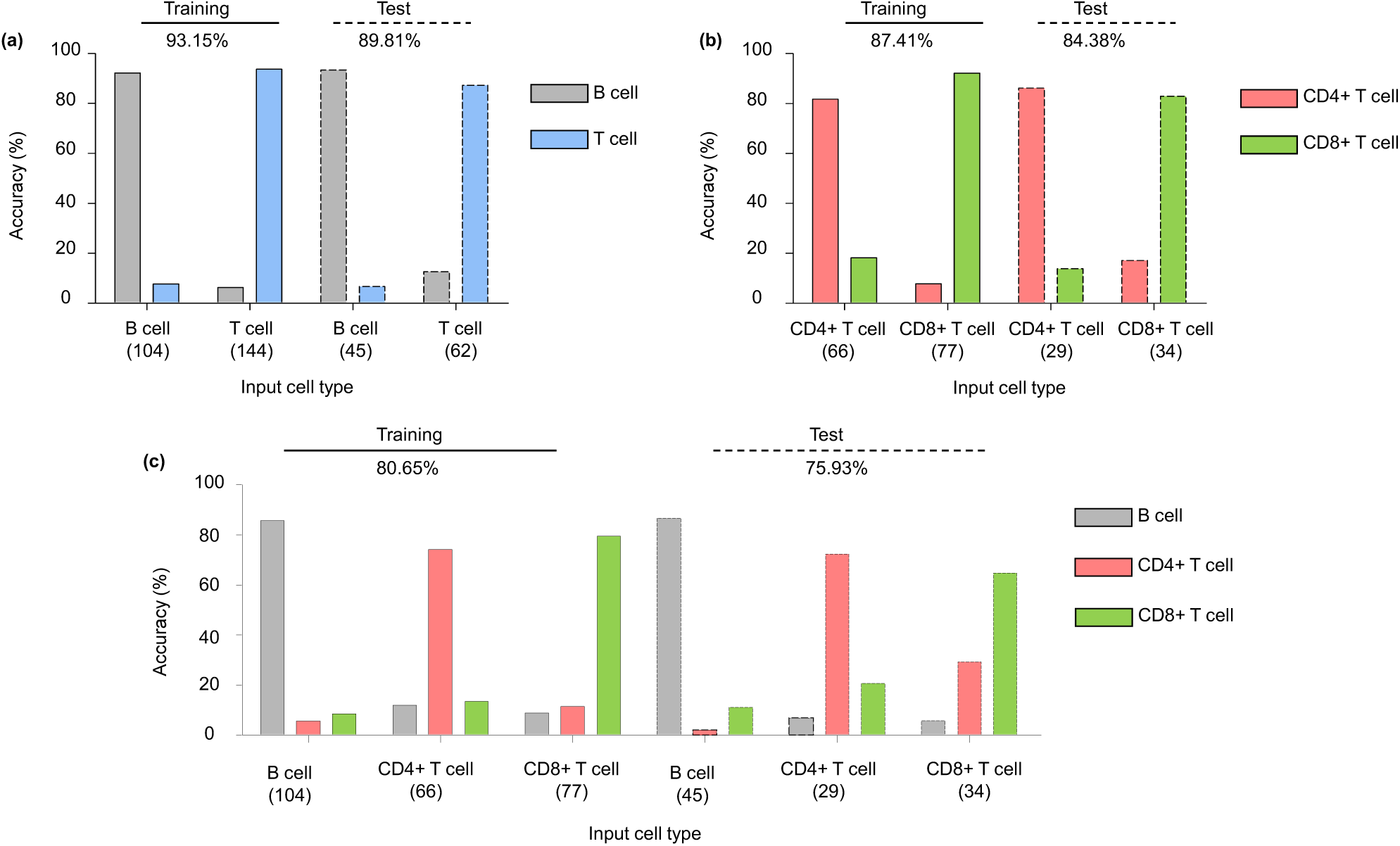
Identification of lymphocyte cell types using a *k*-nearest neighbour algorithm. Identification results of (a) B and T cell types, (b) T cell subtypes (CD4+, CD8+), and (c) all three lymphocyte types. Numbers below each cell type indicate the number of cells used for supervise learning and identification.

**Table 1.**
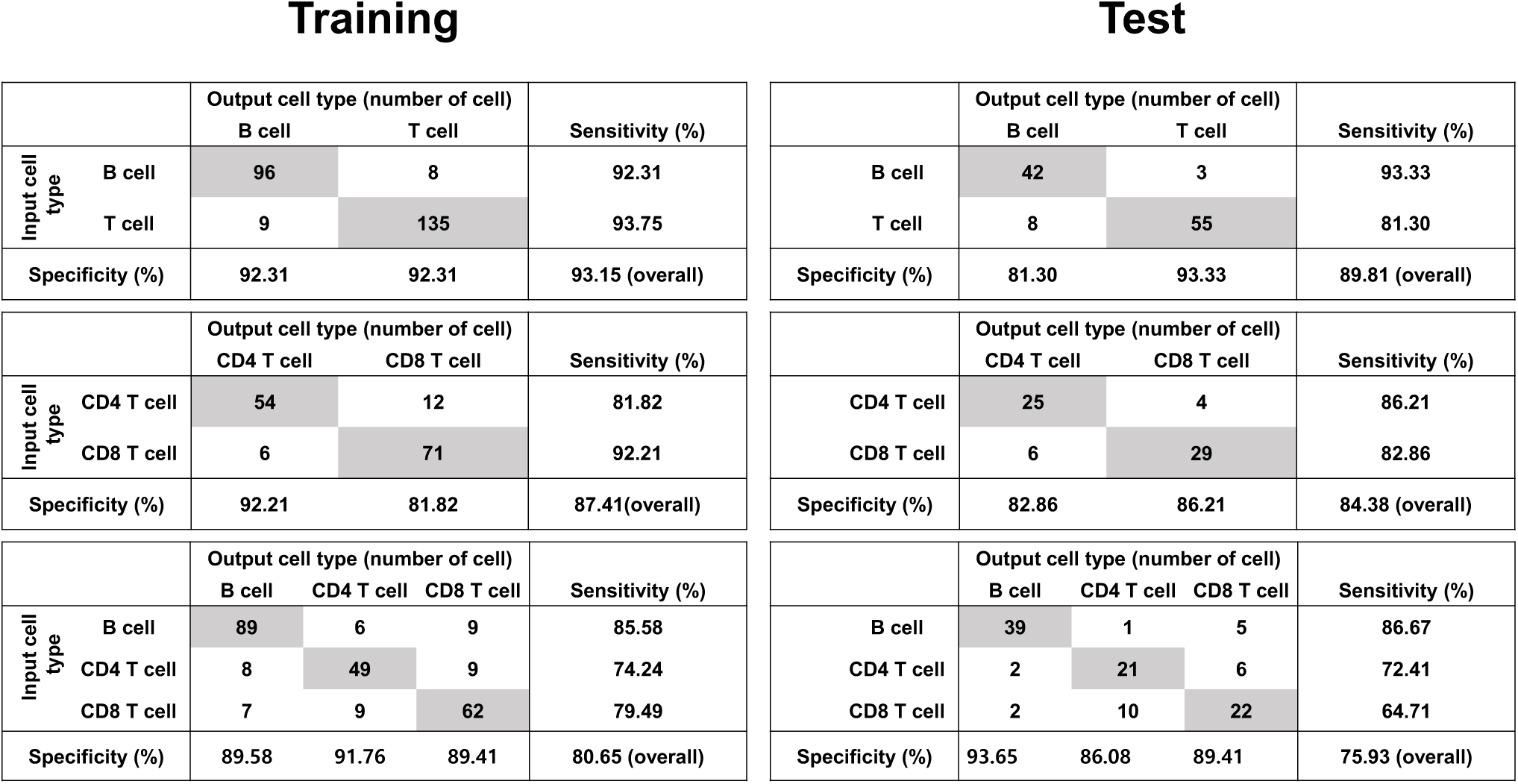
Results of the identification of lymphocyte cell types using a *k*-nearest neighbour (*k* = 4) algorithm.

Training
|  |  | Output cell type (number of cell) |  | Sensitivity (%) |
| --- | --- | --- | --- | --- |
|  |  | B cell | T cell |  |
| Input cell type | B cell | 96 | 8 | 92.31 |
|  | T cell | 9 | 135 | 93.75 |
| Specificity (%) |  | 92.31 | 92.31 | 93.15 (overall) |
|  |  | CD4 T cell | CD8 T cell |  |
| Input cell type | CD4 T cell | 54 | 12 | 81.82 |
|  | CD8 T cell | 6 | 71 | 92.21 |
| Specificity (%) |  | 92.21 | 81.82 | 87.41(overall) |

|  |  | Output cell type (number of cell) |  |  | Sensitivity (%) |
| --- | --- | --- | --- | --- | --- |
|  |  | B cell | CD4 T cell | CD8 T cell |  |
| Input cell type | B cell | 89 | 6 | 9 | 85.58 |
|  | CD4 T cell | 8 | 49 | 9 | 74.24 |
|  | CD8 T cell | 7 | 9 | 62 | 79.49 |
| Specificity (%) |  | 89.58 | 91.76 | 89.41 | 80.65 (overall) |

Test
|  |  | Output cell type (number of cell) |  | Sensitivity (%) |
| --- | --- | --- | --- | --- |
|  |  | B cell | T cell |  |
| B cell |  | 42 | 3 | 93.33 |
| T cell |  | 8 | 55 | 81.30 |
| Specificity (%) |  | 81.30 | 93.33 | 89.81 (overall) |
|  |  | CD4 T cell | CD8 T cell |  |
| CD4 T cell |  | 25 | 4 | 86.21 |
| CD8 T cell |  | 6 | 29 | 82.86 |
| Specificity (%) |  | 82.86 | 86.21 | 84.38 (overall) |

|  |  | Output cell type (number of cell) |  |  | Sensitivity (%) |
| --- | --- | --- | --- | --- | --- |
|  |  | B cell | CD4 T cell | CD8 T cell |  |
| B cell |  | 39 | 1 | 5 | 86.67 |
| CD4 T cell |  | 2 | 21 | 6 | 72.41 |
| CD8 T cell |  | 2 | 10 | 22 | 64.71 |
| Specificity (%) |  | 93.65 | 86.08 | 89.41 | 75.93 (overall) |

## Discussion

We demonstrated label-free identification of lymphocyte cell types at a single cell level using ODT and statistical classification. ODT provides quantitative morphological and biochemical information on the lymphocytes by measuring their 3-D RI distribution. We found that there were significant differences in the quantitative characteristics among the lymphocyte cell population; however, individual lymphocytes in different lymphocyte cell types are indistinguishable due to cell-to-cell variations. To overcome this limitation, the *k*-NN (*k* = 4) algorithm was exploited as a statistical classification method to establish classification models using the quantitative characteristics of the lymphocytes. The optimized classification models can discriminate B and T cells with high accuracy. In addition, the T cell subsets were identified using the *k*-NN (*k* = 4) algorithm with an overall accuracy of over 80%. Moreover, the three types of lymphocytes were identified using the classification model with an overall accuracy of over 75%.

The identification results show that the classification models more precisely discriminate between B and T lymphocytes rather than the T cell subsets, which implies that differences in cellular morphology and biochemical properties between B and T cells are more distinct than those between the CD4+ and CD8+ T cells. The results are consistent with previous knowledge of the lymphocyte-differentiation pathway^48^. B and T lymphocytes originate from hematopoietic stem cells and then mature in different organs. Thus, lymphocytes have identical cellular phenotypes such as one large nucleus and spherical shapes; however, B and T lymphocytes have entirely different cellular functions. Even though our method established the classifiers by optimizing features without biological relevance, the machine learning algorithm found distinct differences in the morphological and biochemical properties among the lymphocyte subtypes.

The present method combines 3-D RI tomography and machine learning, which provides several advantages. First, the present method enables label-free identification of lymphocyte subtypes which cannot be achieved by optical microscopic techniques without using fluorescent methods because of similar phenotypes in lymphocyte subtypes. ODT measures the 3-D RI distribution of the lymphocytes, and machine learning methods find significant differences among the lymphocyte subsets to identify their cell types. Second, the present method has a simple and cost-effective optical setup compared to flow cytometry or other label-free techniques such as Raman spectroscopy. Recently, a 3-D holographic microscope has become commercially available, which simplifies the optical system and reduces the required time for measuring a 3-D RI tomogram by exploiting a digital micromirror device^49^. Thus, the present method can be easily transferred to basic research facilities and clinics. Lastly, there is no limitation in applying the present method to discriminate other types of cells including WBCs, cancer cells, neurons, and glial cells. Because ODT has been widely used to measure various biological samples, the present approach can be readily used to classify various cell types.

There are several points to be improved in future works. The present study proves a proof of concept; however, the overall accuracies for identifying each lymphocyte type should be enhanced. Several classification algorithms including two *k*-NN algorithms (*k* = 4 and *k* = 6), linear discrimination, quadratic discrimination, naïve Bayes, and decision tree were tested, and the *k*-NN (*k* = 4) algorithm showed the best performance for identifying the lymphocyte types (Supplement Fig. 2). However, these classifiers are at the basic level of machine learning methods. Recently, deep learning methods have been introduced and widely sued in various research fields including image recognition^50^, speech recognition^51^, and biomedical research^52-54^. Thus, the identification accuracy could be more improved by exploiting ODT and deep learning methods. In addition, the speed of measuring the 3-D RI tomograms could be improved. For single lymphocyte imaging, we found and measured lymphocytes sparsely placed on a cover glass which limits the speed of the tomogram measurements. We expect that microfluidic approaches might be a solution to increase the speed of the 3-D RI tomograms of lymphocytes and become a practical method for discriminating cell types.

In summary, we envision that ODT combined with machine learning will be a useful tool in biomedical research. ODT quantitatively provides the morphological and biochemical characteristics of samples, and machine learning enables the classification of cell types using the measured quantitative information. The present method can be further used to study immunology, cancer, and neuroscience.

## Methods

### Mice

C57BL/6J mice (gender and age-matched, 6-8 weeks) were purchased from Daehan Biolink (Korea). Animal care and experimental procedures were performed under approval from the Animal Care Committee of KAIST (KA2014-01 and KA2015-03). All the experiments in this study were carried out in accordance with the approved guidelines

### Flow cytometry and Lymphocyte Sorting

White blood cells were isolated from the blood harvested from the heart of mice. Erythrocytes were removed by ACK lysis. Cells were blocked with anti-CD16/32 and then stained for surface molecules. DAPI (4,6-diamidino-2-phenylindole; Roche) was used for dead cell exclusion. Sorting was performed on an Aria II or III system (BD Biosciences) using an 85-μm nozzle or Astrios system (Beckman Coulter) using a 70-μm nozzle. Antibodies for flow cytometry were purchased from BD Biosciences, eBioscience, Biolegend. The antibodies used were CD3ε (clone 17A2), CD4 (GK1.5), CD8α (53-6.7), CD19 (1D3), CD45R (B220, RA3-6B2), NK1.1 (PK136).

### Measurement of 3-D refractive index tomograms

To reconstruct the 3-D RI tomograms of lymphocytes, a Mach-Zehnder interferometric microscope was used^55^ (Fig. 2). A laser beam from a diode-pumped solid-state laser (*λ* = 532 nm, 100 mW, Shanghai Dream Laser Co., Shanghai, China) is split into two arms using a beam splitter. One arm illuminates a sample with various illumination angles ranging from – 60° to 60° in air at the sample plane with respect to the optic axis, which is systematically controlled with a dual-axis galvanomirror (GVS012, Thorlabs, Newton, NJ, USA), and the other is used as a reference beam. The sample is placed between a condenser lens (UPLSAPO Water 60×, the numerical aperture (NA) = 1.2, Olympus, Japan) and an objective lens (PLAPON Oil 100×, NA = 1.4, Olympus, Japan). The diffracted light from the sample is then collected by the objective lens and projected onto the camera plane. At the camera plane, the sample beam interferes with the reference beam, generating spatially modulated holograms, which are then captured by a CMOS camera (1024 PCI, Photron USA Inc., San Diego, CA, USA). For reconstructing a 3-D RI tomogram, a total of 300 holograms of a sample are measured by changing the angle of illuminations which takes less than 1 sec. Then, the optical field information (amplitude and phase) of the measured holograms are retrieved using a field retrieval algorithm based on Fourier transformation^56,57^. From the retrieved multiple amplitude and phase information, a 3-D RI tomogram is reconstructed using an optical diffraction tomography algorithm. An iterative regularization algorithm with a non-negatively constraint was used to fill the missing cone information which results from the limited NA of the condenser and objective lenses^58^. Details on reconstructing 3-D RI tomograms can be found elsewhere^15,59^.

### Quantitative characterization of the structural and biochemical information of the lymphocytes

Quantitative structural and biochemical information of the lymphocytes were calculated from the measured 3-D RI tomograms. To calculate the cellular volume *V* and surface area *S* of a lymphocyte, the voxels of a 3-D RI tomogram of a lymphocyte, which had higher RI values compared to the threshold RI value, were selected. From the selected voxels, the cellular volume and surface area were calculated corresponding to the numbers of voxels and cellular boundaries, respectively. Sphericity, which is a dimensionless parameter and indicates the roundness of a lymphocyte, was obtained from the measured cellular volume and surface area as follows: *Sphericity* = π^*1/3*^·*(6V)^2/3^/S*. The biochemical information (dry mass density and cellular dry mass) were obtained from the RI values due to the linear relation between an RI value and the local concentration of non-aqueous molecules (i.e., proteins, lipids, and nucleic acids inside cells). RI values were converted to the concentration of non-aqueous molecules (mostly proteins) with the following relationship: *n* = *n*_*0*_ +*αC*, where *n* is an RI value of a voxel; *n*_*0*_ is an RI value of the background, and *α* is the refractive index increment (RII) of proteins. Because most proteins have similar RII values, we used a RII value of 0.2 mL/g in this study. The total dry mass of a lymphocyte was calculated by simply integrating the dry mass density over the cellular volume. Details on calculating the quantitative information of a sample can be found elsewhere^23,25^.

### Image processing and statistical analysis

Image processing was performed with Matlab_R2014b and ImageJ. Statistical analysis was done by the GraphPad Prism software. The RI isosurfaces were rendered by commercial software (TomoStudio, Tomocube Inc., Korea).

## ACKNOWLEDGEMENTS

This work was supported by KAIST, Tomocube, and the National Research Foundation of Korea (2015R1A3A2066550, 2014M3C1A3052567, 2014K1A3A1A09063027 to Y.P., 2012M3A9B4027955 to S.K.).

## AUTHOR CONTRIBUTIONS

Y.P. conceived the idea and directed the work. Y.P. and S.K. analysed the data. K.K. designed optics system. M.K. isolated lymphocytes from mice peripheral blood. J.Y. performed the experiments. J.Y. and Y.J established statistical classification model. All authors wrote the manuscript.

